# Ocular findings in Northern Gannets following an outbreak of high pathogenicity avian influenza (H5N1)

**DOI:** 10.64898/2026.04.15.718625

**Authors:** Chloe Fontaine, Ben Blacklock, David Kayes, Josie Parker, Emma Cunningham, Hannah Ravenswater, Jana WE Jeglinski, Elizabeth Mackley, Kirsty A Franklin, Claudia Tapia-Harris, Adrian W Philbey, Liam A Wilson, Mariana Santos, Jude V Lane

**Affiliations:** Hospital for Small Animals, Royal (Dick) School of Veterinary Studies, University of Edinburgh, UK; Roslin Institute, Royal (Dick) School of Veterinary Studies, University of Edinburgh, UK; Institute of Ecology and Evolution, School of Biological Sciences, University of Edinburgh, UK; School of Biological Sciences, University of Edinburgh, UK; Department of Ecoscience, Aarhus University, Roskilde, Denmark.; RSPB Centre for Conservation Science, The Lodge, Sandy, Bedfordshire, UK; Easter Bush Pathology, Royal (Dick) School of Veterinary Studies, University of Edinburgh, UK; Institute of Zoology, Zoological Society of London, Regent’s Park, London, UK

**Keywords:** *Morus bassanus*, avian influenza virus, iridal pigmentation, uveitis, HPAI, black-eye phenotype

## Abstract

**Background:** During 2021–2022, high pathogenicity avian influenza (HPAI) caused mass mortality in wild birds across Europe, with Northern Gannets (*Morus bassanus*) among the most affected. Following the outbreak, unusual alterations in the species’ characteristic pale iris were observed in some individuals.

**Methods:** Opportunistically captured gannets on Bass Rock (n=52), selected to represent a range of iris pigmentation, were examined. Slit-lamp biomicroscopy, indirect ophthalmoscopy, rebound tonometry and photography were performed. Iris pigmentation was classified as normal, mottled or black. Eleven birds underwent avian influenza virus (AIV) serology. Histopathology was performed on two eyes.

**Results:** Abnormal iris pigmentation was found in 74% of adult and immature gannets, with 61% affected bilaterally. Additional signs consistent with uveitis were present in 77% of affected birds. Iris pigmentation abnormalities were positively associated with AIV H5 seropositivity (Fisher’s exact test, P=0.018). Histopathology from affected eyes showed increased melanin deposition and disorganisation, including loss of a distinct anterior layer of melanocytic cells and hypertrophy of melanocytes within the iris stroma.

**Limitations:** Field conditions limited uniform lighting and concurrent serology.

**Conclusions:** Iris pigmentation changes were associated with prior HPAI exposure and frequently accompanied by signs of uveitis, suggesting iris alterations may indicate past infection and potential chronic sequelae.

## INTRODUCTION

Avian influenza viruses (AIVs) are negative-sense, single-stranded RNA viruses belonging to the family *Orthomyxoviridae* and are classified as low or highly pathogenic (1). High pathogenicity avian influenza (HPAI) can produce diverse clinical outcomes across affected species. In some hosts, infection results in rapid multi-organ failure and death with minimal preceding signs, whereas others may exhibit neurological or systemic manifestations (2).

The Northern Gannet (*Morus bassanus*), hereafter gannet, is the largest breeding seabird in the North Atlantic and was one of the most evidently impacted species during the 2021–2022 HPAI H5N1 outbreak (3,4). Until recently, the world’s largest gannet colony was located on Bass Rock, Scotland (Figure 1) (5). The colony experienced a substantial decline following the outbreak, with the number of apparently occupied nest sites decreasing by approximately 43%, from an estimated 81,000 in 2021, prior to the outbreak, to 46,045 in 2024 (6). Despite high mortality, a proportion of remaining gannets in the colony exhibited marked alterations in ocular appearance, with the characteristic pale iris appearing to become partially or completely black (4). This striking phenotypic change is highly unusual and has not been reported in the scientific literature prior to the HPAI outbreak.

**Figure 1:**
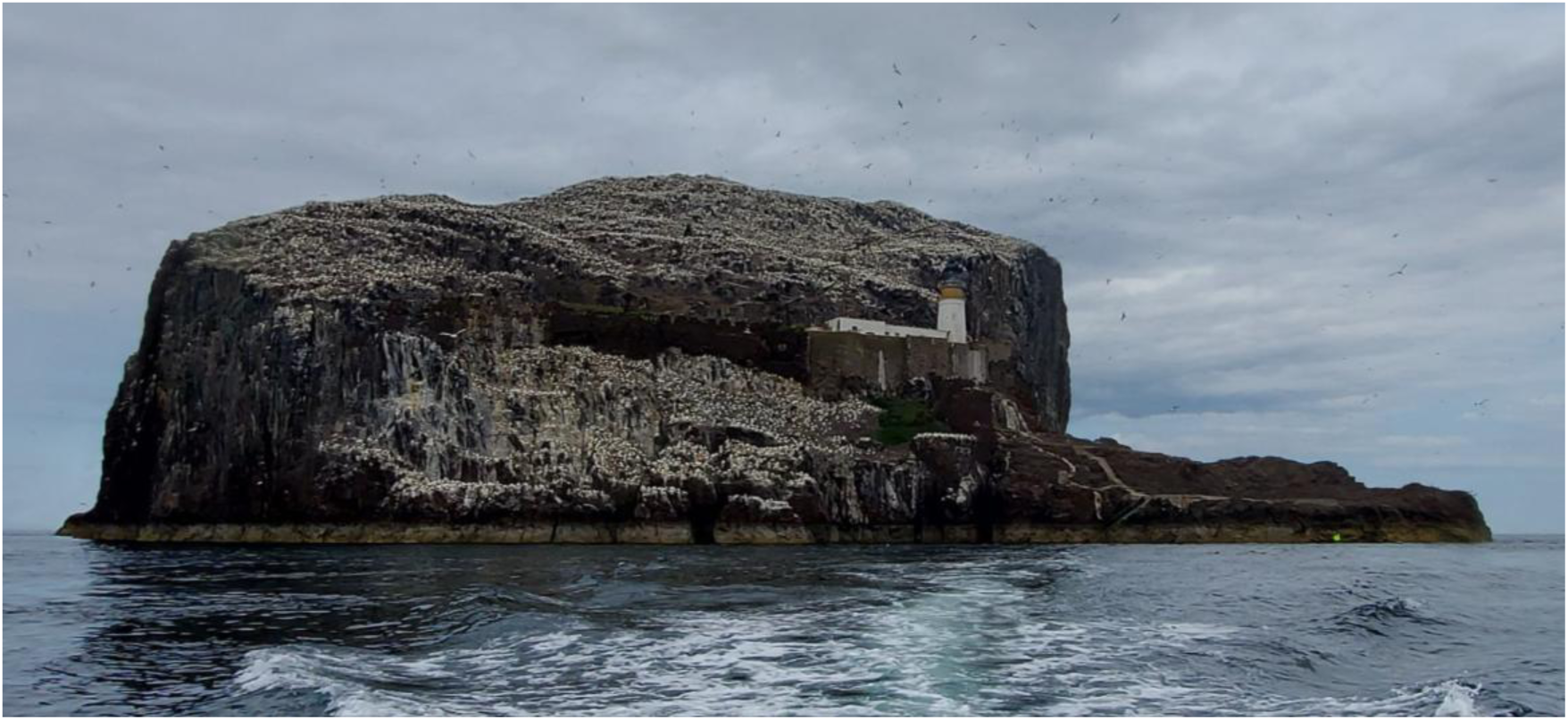
Bass Rock, Scotland.

The aim of this study was to 1) describe the ophthalmic findings observed in gannets, with both normal and affected eyes, following the 2022 HPAI outbreak, 2) investigate the relationship between ocular changes and serological status, and 3) provide histopathological observations of eyes with and without ocular changes.

## MATERIALS AND METHODS

### Ophthalmic examination

Ophthalmic data were collected from gannets on Bass Rock, Scotland. Birds caught under licence using a 6m pole fitted with a noose, for the purposes of ringing and GPS tracking, were examined by a European College of Veterinary Ophthalmologists (ECVO) board-certified veterinary ophthalmologist or an ECVO ophthalmology resident during four trips to the island in April, July and August 2025. Birds were opportunistically selected to include a range of eye morphologies including normal eyes. Each bird, unless already ringed, was fitted with a metal British Trust for Ornithology (BTO) leg-ring and a coloured acrylic leg-ring engraved with a unique 4-digit alphanumeric code for future identification.

The following variables were recorded for each adult and immature bird: 1) intraocular pressure (IOP), measured using rebound tonometry (p setting on TonoVet, iCare, Helsinki, Finland); 2) slit-lamp biomicroscopic findings (SL-17, Kowa Optimed, Japan); 3) presence or absence of aqueous flare; 4) iris colour classification; 5) photographs of both eyes, captured using a digital single-lens reflex (DSLR) camera equipped with a lens and flash. Birds were restrained using a custom-made jacket and the head held stationary by a fieldworker experienced in handling gannets. Bird handlers wore safety glasses, cut-resistant gloves and arm sleeves to protect against the birds’ sharp beak. A firm grip on the neck and head, and close control of the beak without clamping it shut to allow the bird to breathe freely, was required throughout the examination, to ensure the safety of the handler and examiner, and to minimise bird movement (Figure 2).

**Figure 2:**
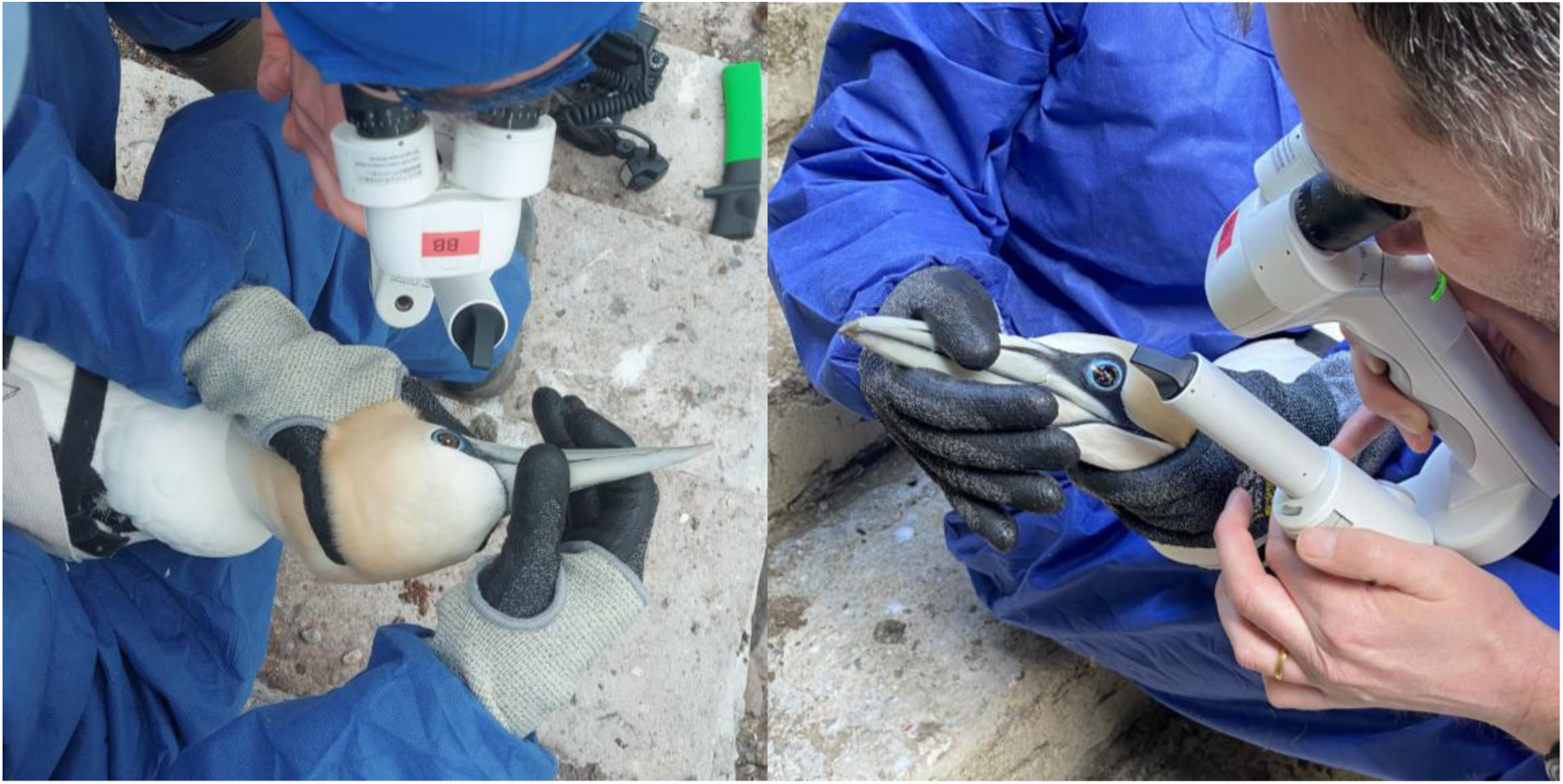
Control of the head and beak for the examination was maintained through a firm hold at the base of the skull and a hold of the beak, without completely closing it. This ensured safety for both handler and examiner, as well as minimising bird movement but enabling the bird to breathe freely. The people in these photographs are authors of this manuscript.

Birds from three age classes, determined by plumage characteristics, were examined: adults (≥5 years; n = 38), immatures (third and fourth calendar years; n = 4), and pre-fledging juveniles (hatched in 2025; n = 10) (Figure 3). Given the similar ocular appearance of adults and immatures, these age classes were combined for statistical analysis. Juveniles were included to provide descriptive baseline ophthalmic observations for this species. As juveniles could not exhibit post-outbreak changes, they were not included in any statistical analyses. Based on photographic assessment, iris pigmentation was categorised independently by two observers (ECVO-boarded ophthalmologists) who were blinded to serological status. Pigmentation was classified as normal, mottled or black. Eyes in which less than 50% of the iris surface was subjectively judged to be affected by pigmentation were classified as mottled, whereas eyes in which 50% or more of the iris surface was pigmented were classified as black. Only birds with complete ophthalmic examination data were included in the statistical analyses.

**Figure 3.**
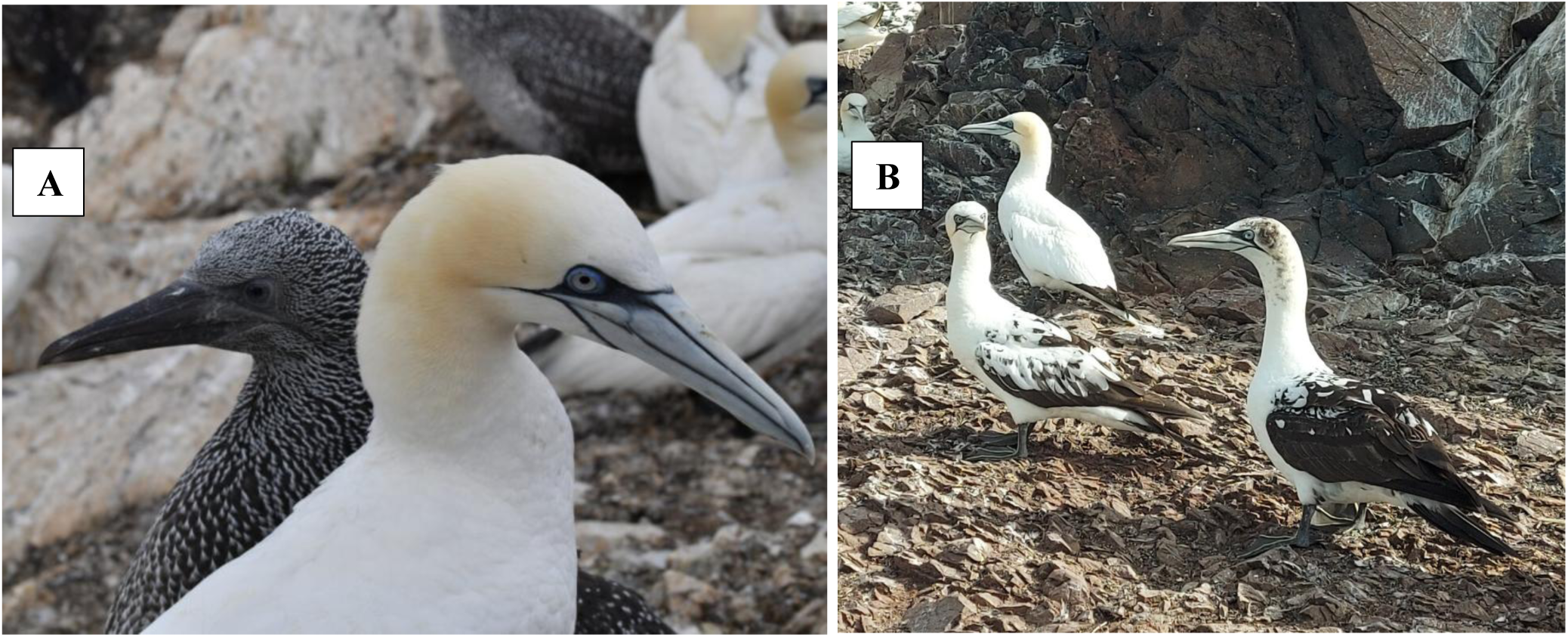
**(A)** Juvenile (dark plumage) gannet next to an adult (**w**hite plumage). **(B)** Immature gannets.

### Serological analysis

#### Blood sample collection

To investigate the link between previous exposure to avian influenza and ocular changes, blood samples were analysed from 11 gannets that also underwent an ophthalmic examination: nine birds with iridal pigment changes and two birds with normal eyes (Table 1). Up to 1.5ml of blood was withdrawn from the medial metatarsal vein using a 25-gauge needle and syringe, placed into a heparinised Eppendorf tube (∼8 µl heparin). Samples were centrifuged at 3000rpm for 16 minutes, after which plasma was separated from red blood cells and kept refrigerated until freezing at −80°C. Sampling was carried out in 2024 (n=5) and 2025 (n=6) under a UK Home Office licence, and in accordance with the Animals (Scientific Procedures) Act 1986.

**Table 1:**
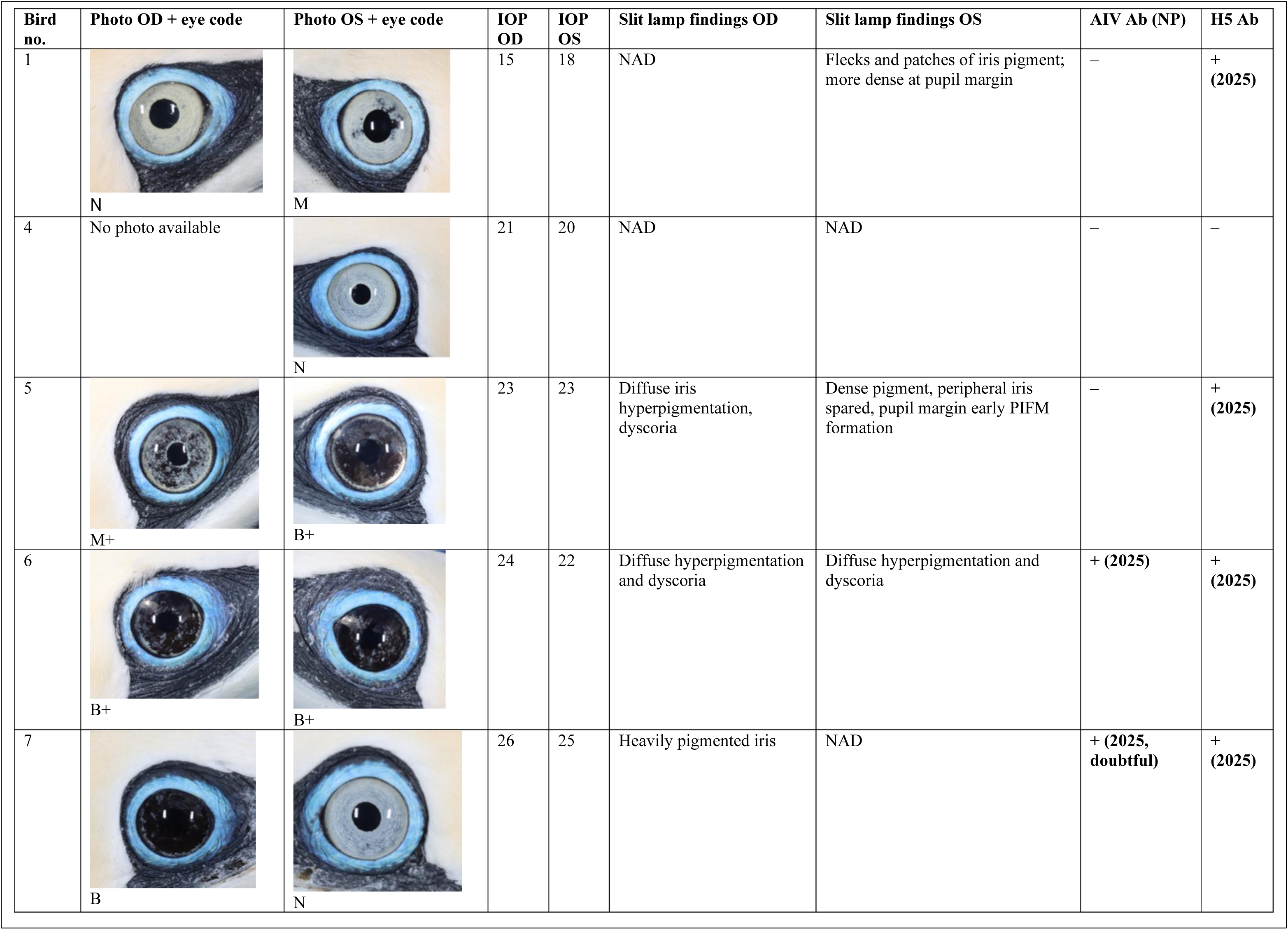
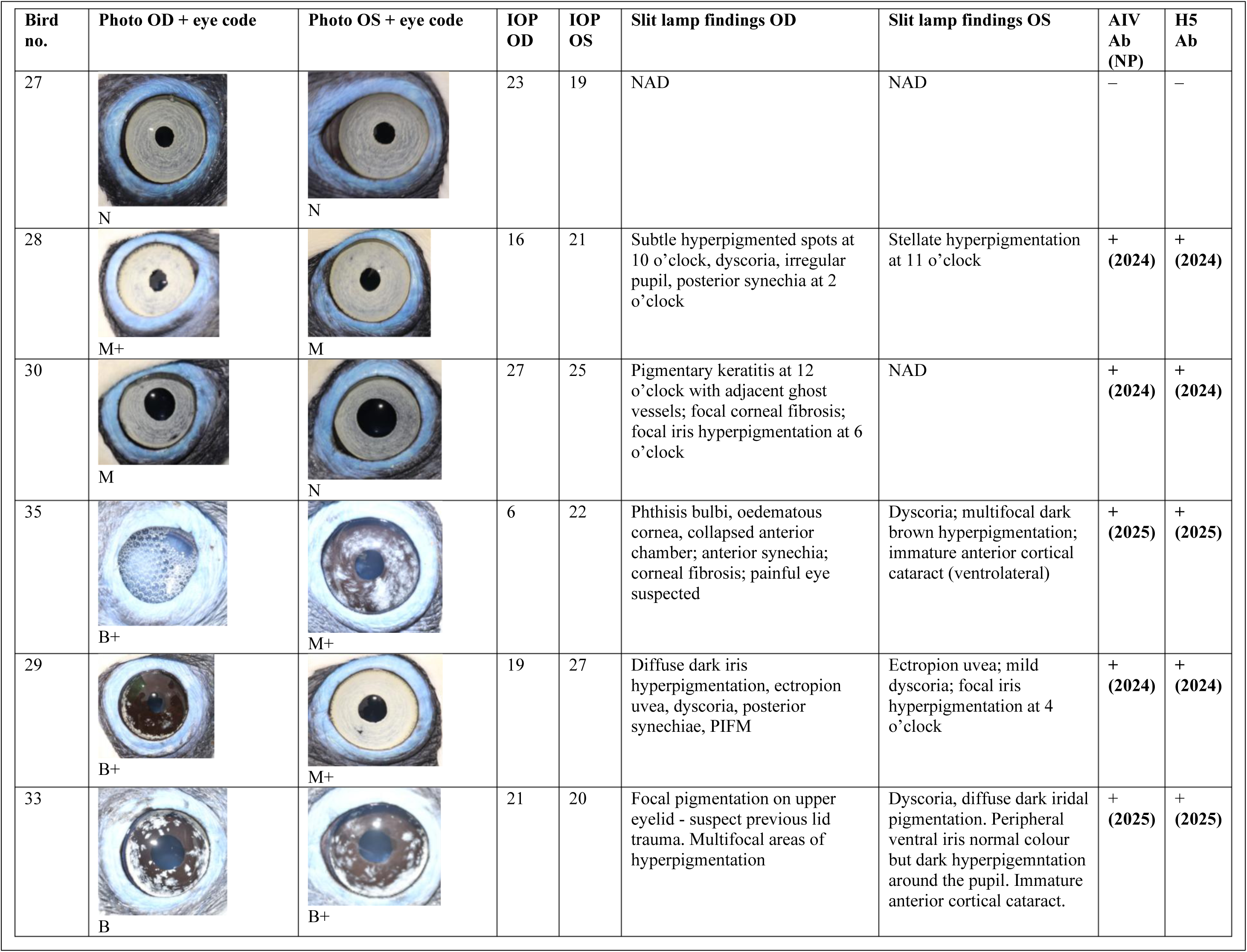
Ophthalmic findings and avian influenza virus (AIV) serology results for the 11 adult gannets included in analysis linking H5 seropositivity and iris pigmentation. Iris pigmentation is classified as normal (N), mottled (M; <50% of iris surface affected), or black (B; ≥50% of iris surface affected), with a “+” suffix indicating additional signs consistent with uveitis. Right eye (OD) and left eye (OS) are shown separately. IOP denotes intraocular pressure, NAD indicates no abnormalities detected, and IAV Ab (NP) denotes avian influenza virus nucleoprotein antibodies.

**Table 2:** Frequency of iridal pigmentation codes for the right (OD) and left (OS) eyes of 42 adult gannets. B: Black iris, M: Hyperpigmented iris in a mottled pattern, N: Normal pale-grey coloured iris, +: With signs associated with uveitis.

| Code | OD | OS |
| --- | --- | --- |
| **B** | 3 | 2 |
| **B+** | 9 | 9 |
| **M** | 5 | 6 |
| **M+** | 9 | 7 |
| **N** | 16 | 17 |
| **N+** | 0 | 1 |
| Total | **42** | **42** |

#### General screening for antibodies to AIV nucleoprotein (NP ELISA)

To test for previous exposure to any AIV, plasma from 11 individuals was assayed for antibodies against the nucleoprotein (NP) of avian influenza, which is conserved across all subtypes H1-H16 and N1-N9. A commercially available competitive ELISA kit (ID Screen® Competitive Multispecies Avian Influenza kit, IDvet, Grabels, France) was used in accordance with the manufacturer’s instructions for avian plasma and serum samples, diluted 1:5. Samples were classified as positive or negative for avian influenza antibodies according to the manufacturer-defined thresholds, based on the sample-to-negative control ratio expressed as a percentage (S/N%). Samples with an S/N% <45% were considered positive, those with an S/N% >50% were considered negative, and values between 45% and 50% were classified as doubtful. All samples were tested in duplicate to ensure repeatability of results.

#### H5 subtype specific antibodies (H5 ELISA)

Following general screening for avian influenza antibodies, NP positive samples were tested for the presence of haemagglutinin H5 specific antibodies. Additionally, one “doubtful” sample and two samples that were NP negative but came from birds that had iridal hyperpigmentation were also tested. A commercially available competitive ELISA (ID Screen Influenza H5 Antibody Competition 3.0 Multi-species ELISA, ID Vet, Montpellier, France), was used. The assay was performed in accordance with the manufacturer’s instructions, following the protocol specified for duck serum, with samples diluted 1:5.

Results were interpreted using the manufacturer-defined S/N% thresholds: samples with S/N% <50% were classified as positive, those between 50% and 60% as doubtful and those >60% as negative. All samples were tested in duplicate to ensure repeatability.

#### Link between seropositivity to H5 antigens and iris pigmentation

For analyses investigating the link between seropositivity to H5 antigens and iris pigmentation, only birds (n=11) with paired ophthalmic and serological data were included. Associations between iridal pigment change and serological status were assessed using Fisher’s exact test, with differences considered significant at P < 0.05. Data analysis was performed using a commercially available statistical software JASP Team (2025) JASP (Version 0.19.3).

#### Licencing and Permissions

Permission to conduct work on Bass Rock was granted by NatureScot (Licence 288302). Capture and ringing were performed using BTO-approved methods and ringing permits. Blood sampling was undertaken by licensed personnel under UK Home Office authority (Project Licence no. PEAE7342F;). Access to Bass Rock for scientific purposes was permitted by the landowner.

All ophthalmic examination procedures were performed as part of separate projects requiring gannet capture for ringing and GPS tracking with no additional intervention or disturbance to the birds. The study was reviewed and authorised by the Royal (Dick) School of Veterinary Studies Veterinary Ethical Review Committee (VERC-195.25).

#### Histopathology

Enucleated pairs of eyes were obtained from two gannets euthanased by veterinary surgeons on welfare grounds: one juvenile with normal eyes, and one adult with unilateral iridal pigmentary changes and blindness. Specimens were fixed in 10% neutral buffered formalin and processed routinely into formalin fixed paraffin embedded blocks, sectioned at 4µm thickness and stained with haematoxylin and eosin for examination. No serological analysis was performed on these two birds.

## RESULTS

### Ophthalmic examination

Ophthalmic examinations were performed on 52 gannets, of which 38 were classified as adult, four as immature, and 10 as juvenile (Figure 3). The eyes were positioned laterally, on either side of the beak, but oriented slightly forward and downwards (Figure 4A). The menace response and dazzle reflex were equivocal in all birds during neuro-ophthalmic examination; during restraint, the globe movement and third eyelid excursions across the cornea were continuous, making both a menace response and dazzle reflex impossible to accurately determine (Video l). Pupils were circular or ovoid in all individuals, with no abnormal ocular findings; pupil size fluctuated continuously throughout the examination period.

**Figure 4.**
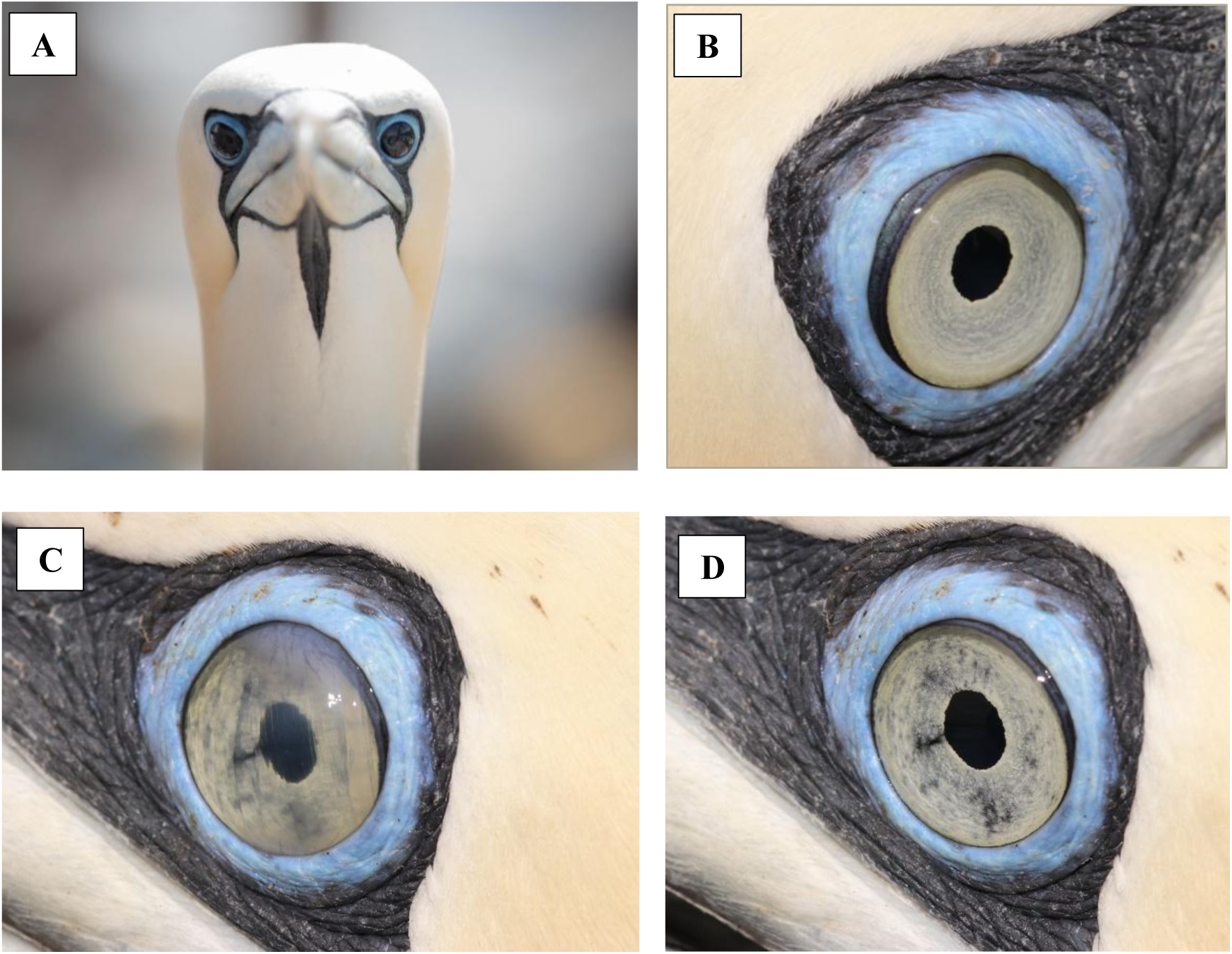
**(A)** Eyes positioned laterally on either side of the beak but oriented slightly forward, and downwards. Right and left darkly pigmented iris in a Northern gannet, categorised as B (Black). **(B)** Characteristic, non-mobile cobalt blue eyelids of a gannet. **(C)** Highly mobile nictitating membrane of a gannet. **(D)** Retracted nictitating membrane (non-visible). Mottled iris pigmentation.

All adult and immature birds had characteristic, non-mobile cobalt blue eyelids (Figure 4B). The nictitating membrane was transparent, highly mobile and moved from nasal to temporal positions (Figure 4C&4D). The eyelids were fixed, and voluntary protrusion of the third eyelid functioned as the equivalent of a blink. Juveniles had a variably brown iris, transitioning to the pale grey adult colouration during the first year of life (Figure 5) (3).

**Figure 5:**
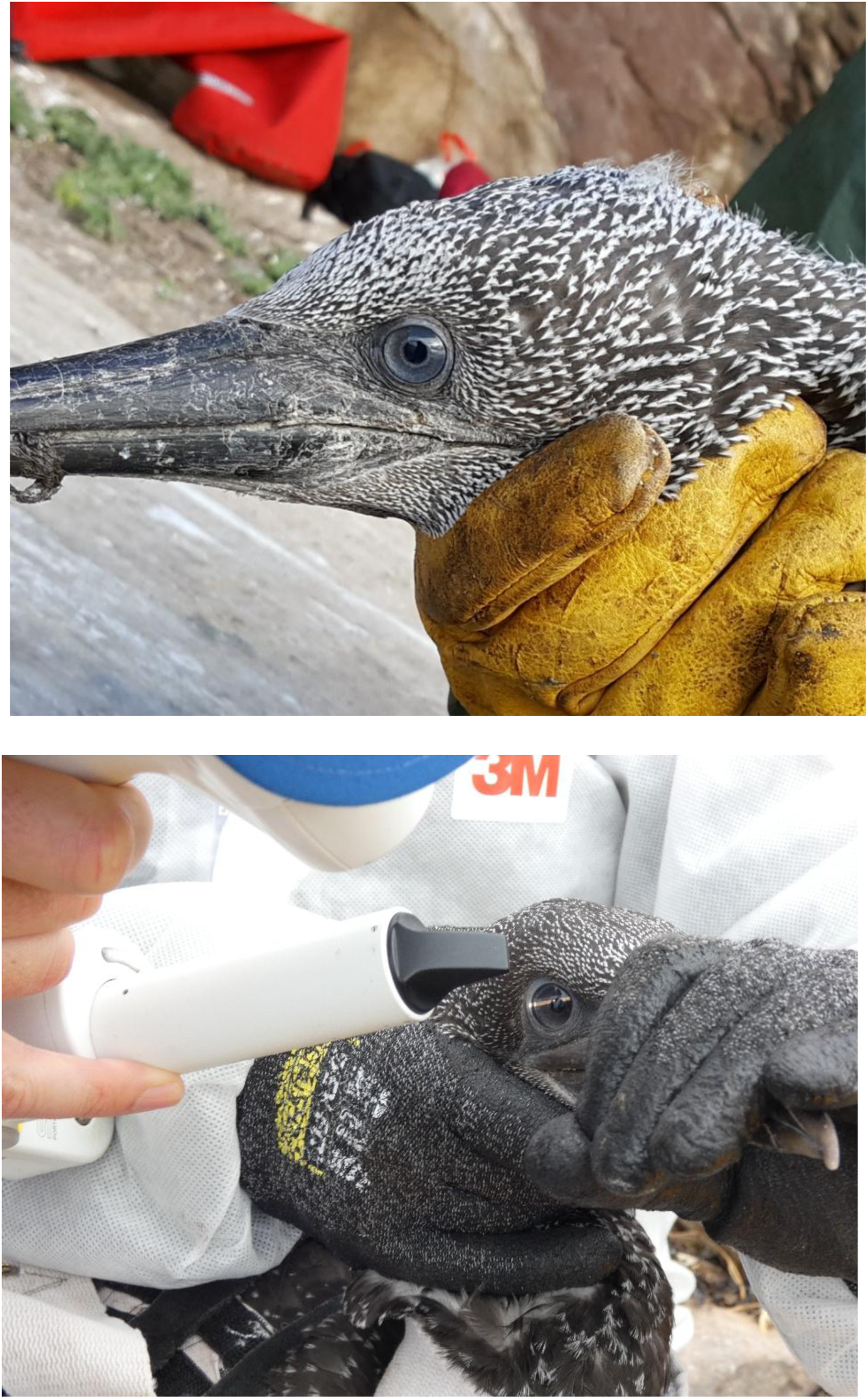
Juvenile eyes with a variably brown iris

### Iridal pigmentation

Among the gannets selected for this study, 26.2% (11/42) had bilaterally normal eyes, while 73.8% (31/42) exhibited abnormal iris pigmentation in at least one eye. All birds with abnormal iris pigmentation were adults that displayed a spectrum of iris changes, ranging from focal mottling (Figure 6A) to complete pigmentation, appearing black in some individuals (Figure 6B). Abnormal iris pigmentation affected 61.3% (19/31) of birds bilaterally, while 38.7% (12/31) were affected unilaterally. Of the 42 pairs of eyes examined, a mottled iris was observed in 14 OD (right eye) and 13 OS (left eye), while a black iris was present in 12 OD and 11 OS.

**Figure 6.**
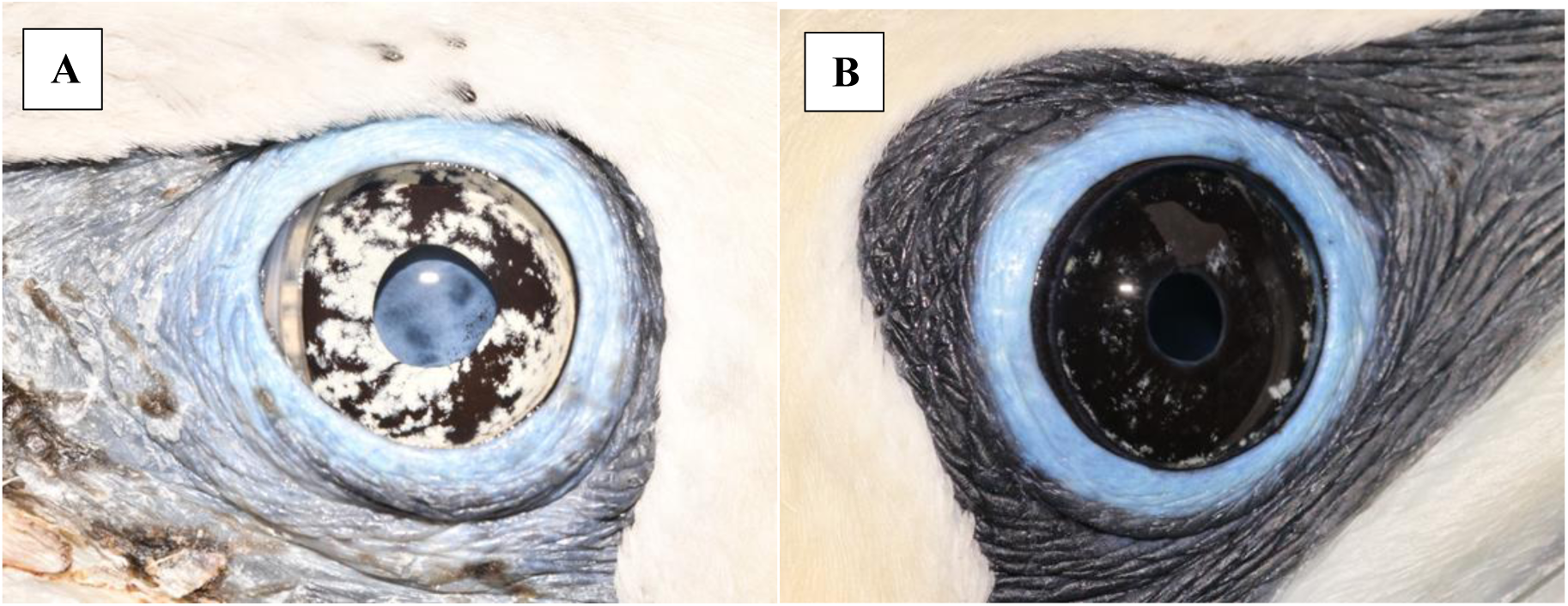
**(A)** Adult eye with mottled iridal hyperpigmentation. Dyscoria due to posterior synechiae at the lateral pupil margin, small spots of pigmentation on the anterior lens capsule, hypermature cataract, posterior lens subluxation noted on slit-lamp examination. **(B)** Adult eye with a black iris. Diffuse dark iris hyperpigmentation, ectropion uvea, dyscoria, posterior synechiae, PIFM observed on slit-lamp examination.

A pale grey (normal) iris was observed in 16/42 OD and 18/42 OS, in which a grey circular pattern of pigmentation (ridges) followed the iris musculature (Figure 7). The ophthalmic examination of the 10 juvenile birds did not reveal any signs of abnormal pigmentation.

**Figure 7:**
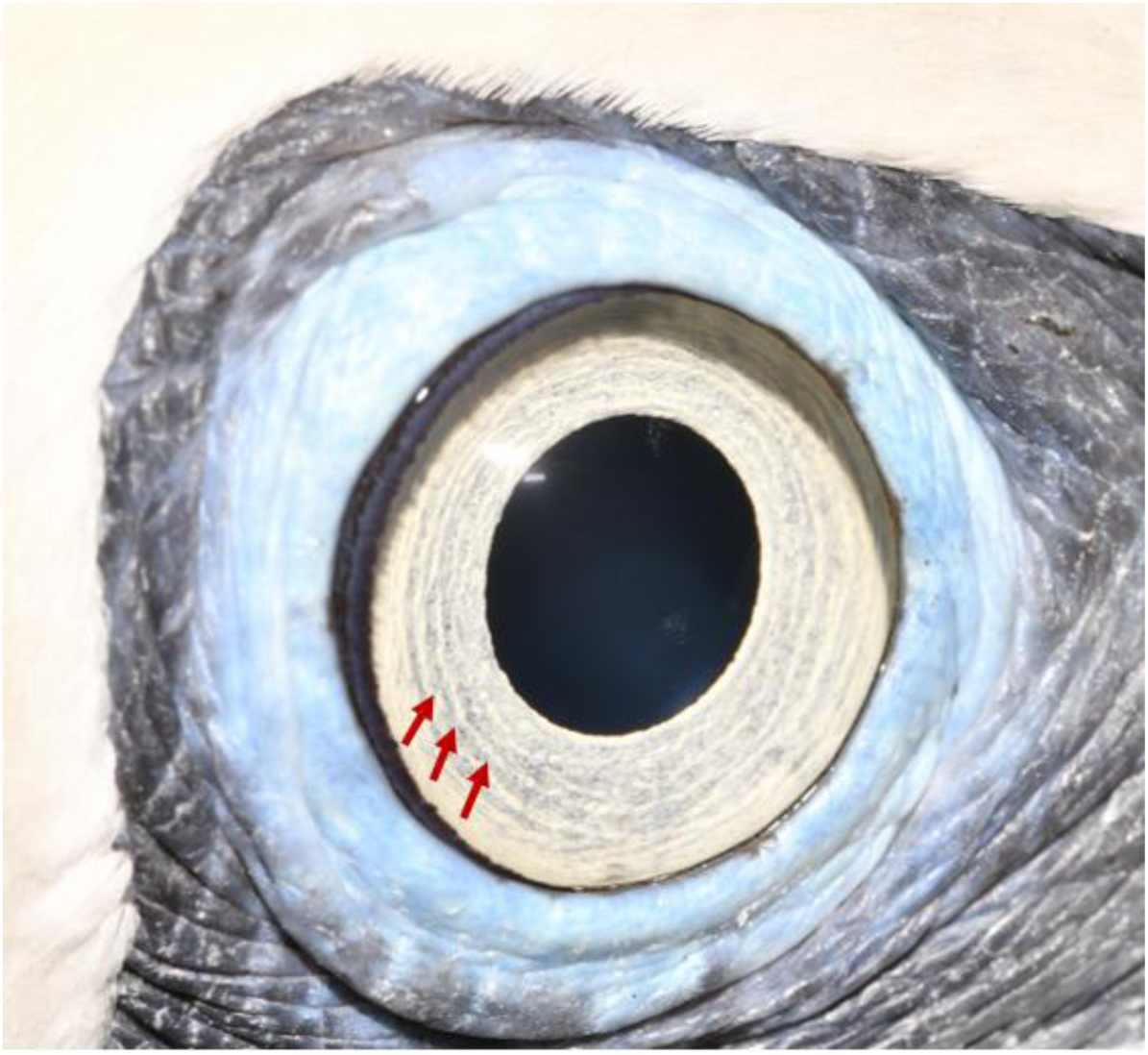
Grey circular pattern of pigmentation (ridges: arrows) following the iris musculature.

### Uveitis

Additional ophthalmic findings associated with uveitis (denoted with a “+” symbol in Table 1) were identified in 77% (24/31) of birds with iridal pigmentation and occurred almost exclusively in eyes with pigmentary change; one bird exhibited dyscoria in a normal left eye but had a mottled right iris. Among eyes exhibiting iris pigmentation (n = 50), dyscoria was the most frequent finding, present in 56% (28/50; 14 OD, 14 OS). Pre-iridal fibrovascular membranes or intraocular fibrin were observed in 16% (8/50; 5 OD, 3 OS), ectropion uvea in 14% (7/50; 4 OD, 3 OS), and anterior or posterior synechiae in 14% (7/50; 6 OD, 1 OS).

Cataract was present in 12% (6/50; 1 OD, 5 OS) of affected eyes (Figure 6A,B).

Phthisis bulbi was identified in one eye of a single bird (2%, 1/50; OD); this bird also had an oedematous cornea, a shallow anterior chamber, and an increased rate of third eyelid excursion resulting in accumulation of air droplets (bubbles) within the tear film (Figure 8). Marked unilateral aqueous humour flare was detected in one bird (2%, 1/50; OD) in April 2025; this bird was subsequently found dead in Sweden (Kattegat) on 5 November 2025.

**Figure 8:**
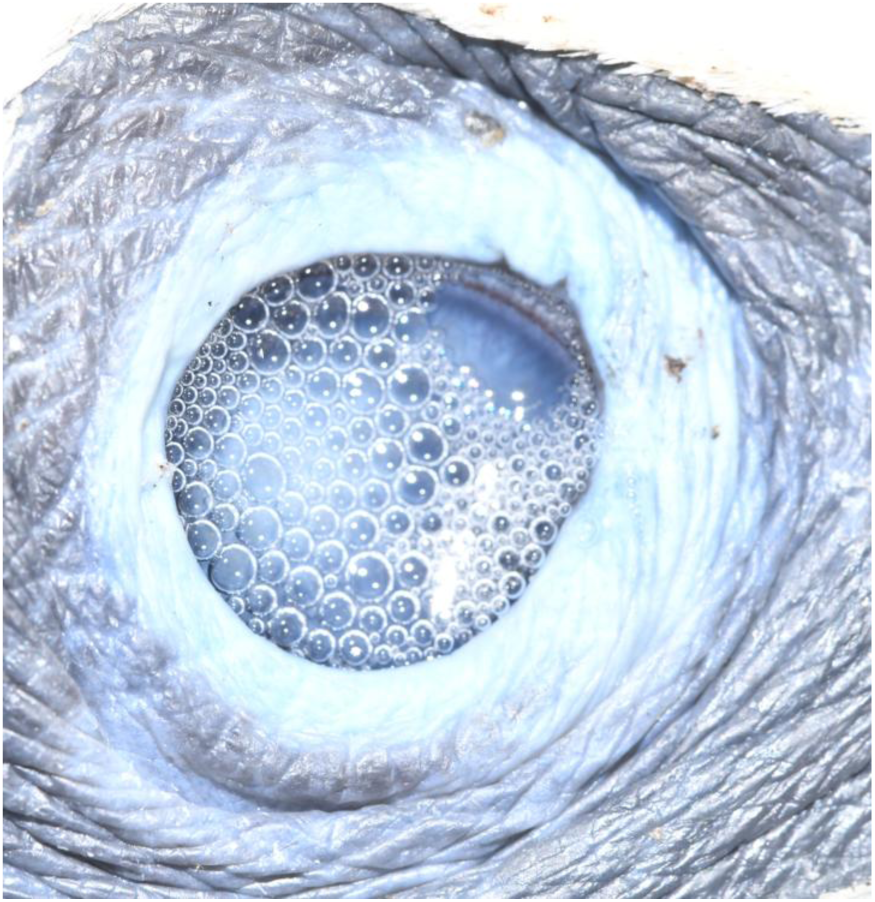
Phthisical right globe with an increased rate of third eyelid protrusion, resulting in a bubbly tear film. Corneal fibrosis is visible ventrolaterally on slit-lamp examination. This individual also exhibited an oedematous cornea, a shallow/collapsed anterior chamber, and anterior synechia.

Among adult gannets with abnormal pigmentation, corneal abnormalities were detected in 6% (3/50; all OD) of affected eyes, including vascular keratitis, pigmentary keratitis, corneal ghost vessels, corneal fibrosis, and corneal oedema; two also exhibited additional signs associated with uveitis.

Mean intraocular pressure (IOP) in adult and immature gannets was 21.9 ± 4.3 mmHg. There was no statistically significant difference in intraocular pressure between pigmented eyes (21.3 ± 4.7 mmHg) and non-pigmented eyes (22.6 ± 3.4 mmHg) (Mann–Whitney U = 910.0, P = 0.38). Ophthalmic examination of all juvenile birds was unremarkable (10/10 normal).

### Serological analysis

Of the 11 individuals tested, 82% (9/11) were positive for H5 antibodies, with no other subtypes detected. All H5-positive birds showed iris pigment change, either mottled or black. The two adult birds that were seronegative for H5 had no iris pigment change; Fisher’s exact test demonstrated a significant positive association between iridal pigment change and AIV H5 seropositivity (P = 0.018) (Table 3).

**Table 3:**
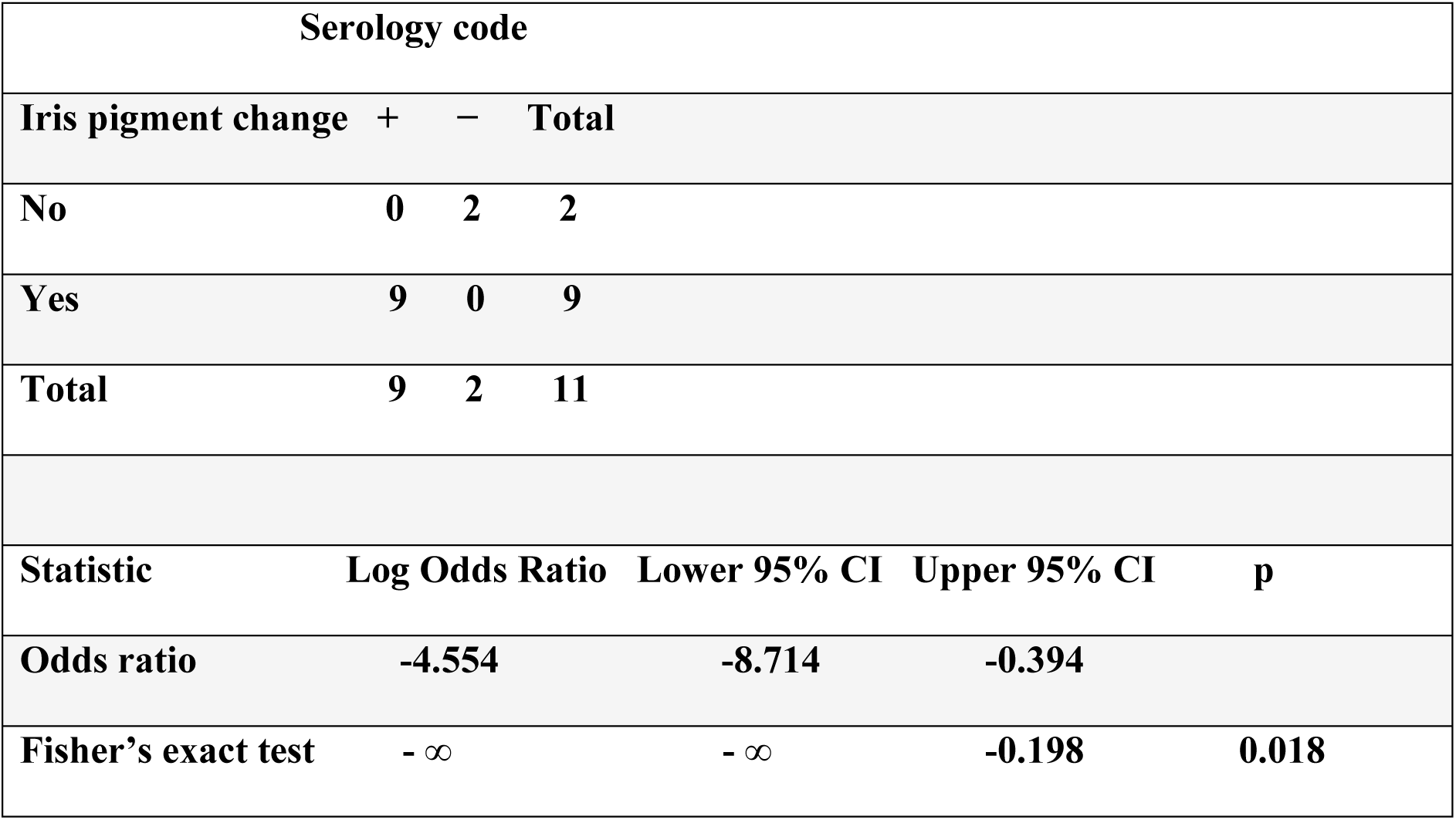
Iridal pigment change in relation to avian influenza virus (AIV) H5 antibodies. Fisher’s exact test shows a significant association between pigment change and AIV H5 seropositivity. “+” indicates seropositivity to H5 antigens, “–” indicates seronegativity. N denotes no iridal pigmentary changes, and Y denotes the presence of iridal pigmentary changes in 11 birds. CI: Confidence interval.

### Histopathology

#### Juvenile eyes

Histopathological examination of the two healthy eyes from the juvenile gannet (Figure 9A) with light brown irises showed mild vascularisation and melanin pigmentation at the corneoscleral junction (limbus), changes considered to be normal findings. The cornea and lens were unremarkable. The iridocorneal angle was open and notably wide compared with that of domestic mammals (e.g. dogs) and birds (e.g. chickens). The anterior border layer of the iris consisted of four to five layers of pigmented cells, overlain by a loose layer of polygonal cells with prominent, finely granular, cytoplasmic pigmentation (presumed melanin) (Figure 9B). The iris stroma was moderately interspersed by melanocytic cells. The iris margin was heavily pigmented, with the posterior iris epithelium extending anteriorly to form a prominent pupillary projection (‘ruff’) (Figure 9B Inset). The ciliary body was composed of a bilayered variably pigmented epithelium, as seen in domestic animals. The uvea otherwise was unremarkable as were the optic disc, optic nerve and pecten. Brucke’s muscle was moderately prominent deep to the scleral ossicles, extended posteriorly to the ciliary body and merged with more anteriorly orientated, fine, skeletal muscle fibres extending anteriorly to the limbus (presumed Crampton’s muscle). The sclera contained scleral ossicles, or bands of bone and cartilage, a normal feature in avian species. The periocular musculature and conjunctiva were unremarkable.

**Figure 9:**
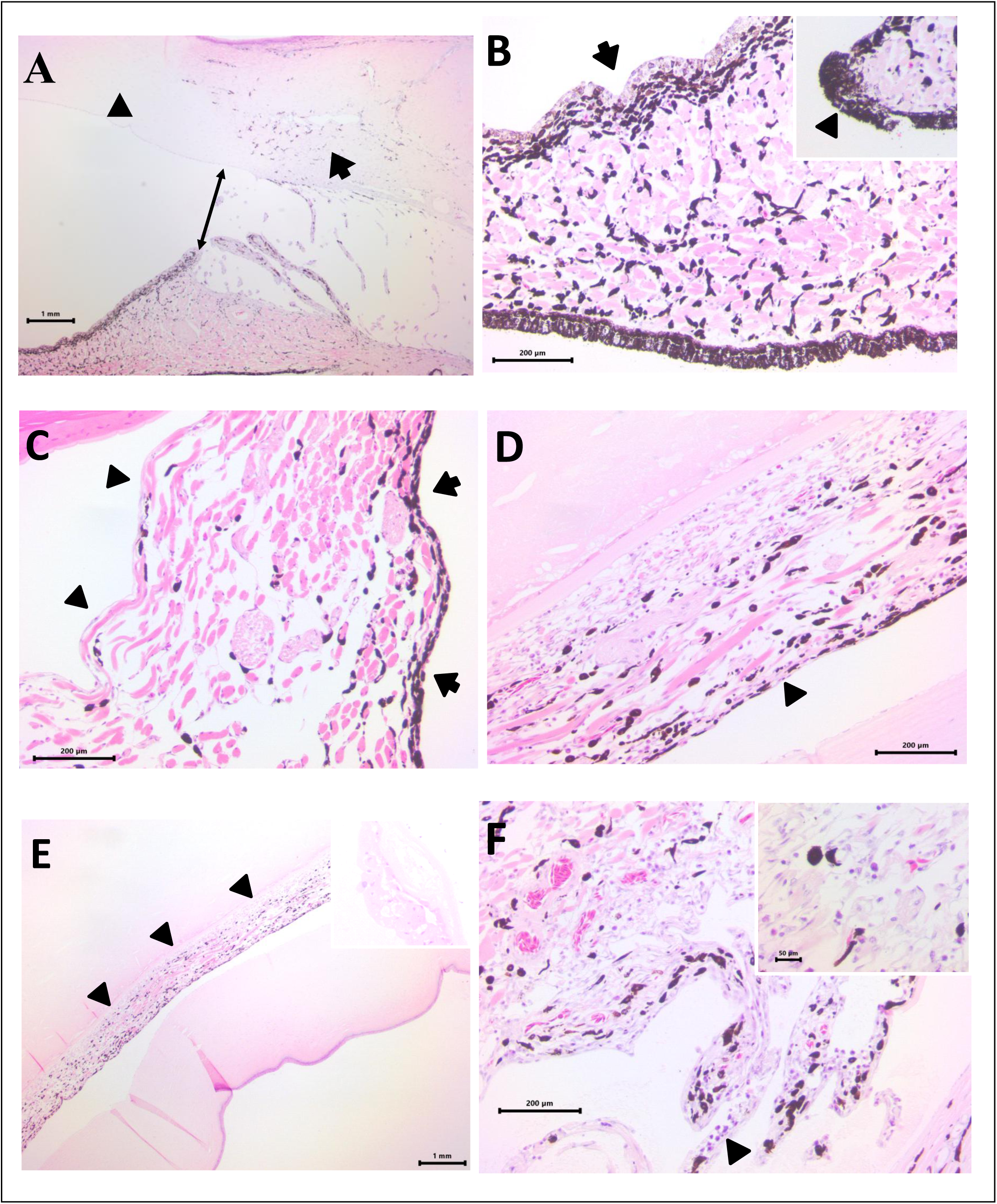
Photomicrographs of gannet eyes **(A)** Normal juvenile gannet eye demonstrating the wide and open filtration angle (double arrow) alongside the scattered pigmentation of the limbus (arrow) with minimal extension into the adjacent cornea (arrowhead). HE, scale bar 1mm. **(B)** Normal juvenile gannet iris demonstrating the distinct and relatively uniform anterior layer of up to four or five pigmented cells (arrow) alongside scattered pigmented cells throughout the iris and a posterior layer of columnar pigmented cells. Inset: At the medial aspect of the pupil, there is a papillary ruff characterised by anterior extension of pigmented cells. HE, scale bar 200µm. **(C)** Clinically normal adult iris demonstrating the loss of the anterior border layer of polygonal, finely granular pigmented cells and loss of pigmented cells in the posterior iris epithelium. HE, scale bar 200µm. **(D)** Adult gannet eye with clinical uveitis. There is marked loss of layering and pigmentation on the anterior aspect of the iris (arrowhead) with subjective hypertrophy of pigmented cells throughout the iris stroma and small numbers of lymphocytes and plasma cells scattered throughout the iris. HE, scale bar 200µm. **(E)** Adult gannet eye with clinical uveitis. There is marked collapse of the anterior chamber with adhesion of the posterior aspect of the iris to the anterior surface of the lens capsule indicative of a posterior synechiae (arrowheads), of which the lens is characterised by a mostly homogenous and lighlty eosinophilic, collapsed stroma. Inset: Scattered remnant bladder cells are present at the periphery of the collapsed lens. HE, scale bar 1mm. **(F)** Adult gannet eye with clinical uveitis, demonstrating extension of the lymphocytic and plasmocytic inflammation (arrowheads) to the adjacent drainage angle. Inset: Higher magnification of uveal inflammation. HE, scale bar 200µm.

#### Adult eye

On clinical examination, the left eye of the adult gannet was visual and appeared normal, whereas the right eye was non-visual with a completely pigmented (black) iris and an aqueous flare, consistent with uveitis.

Histopathological examination of the left (clinically unaffected) eye revealed no significant pathological abnormalities, with an intact cornea, mild vascularisation, and melanin pigmentation at the limbus, considered to be within normal limits and similar to the features described above in the juvenile’s eyes. Brucke’s and Crampton’s muscle were also similar to those described in the juvenile eyes.

On the anterior border layer of the iris, the prominent layer of polygonal cells with granular pigmentation seen in the juvenile eye was absent, and the stroma contained fewer pigmented melanocytes than in the healthy juvenile eye (Figure 9C). The posterior iris epithelium was small, flattened, and indistinct.

In contrast, the right eye demonstrated marked pathological changes consistent with chronic uveitis, including diffuse infiltration of the uvea and pecten by moderate numbers of lymphocytes and plasma cells, with loose perivascular lymphocytic aggregates within the ciliary body. Mild, diffuse oedema was also present throughout the uvea. Interestingly, the iris stroma was similarly sparsely pigmented to the contralateral eye, with similar loss of the anterior layer of polygonal cells with granular pigment and an indistinct posterior iris epithelium. The pigmented cells remaining on the anterior aspect were however diffusely flattened, loosely arranged and separated whilst the pigmented cells throughout the iris stroma were subjectively slightly plump and enlarged (Figure 9D). The cornea, including the epithelium, stroma and Descemet’s membrane, was intact, although moderate vascularisation and melanin pigmentation were present at the corneoscleral junction (limbus), considered to be within normal limits. The lens showed severe degeneration and partial collapse, with liquefaction of lens fibres and replacement by floccular eosinophilic granular material alongside multifocal clusters of peripheral bladder cells but with retention of the lens capsule (Figure 9E Inset), consistent with advanced cataract formation. A few subcapsular lens epithelial cells remained, interpreted as residual anterior lens epithelial cells. The posterior surface of the iris was adherent to the anterior lens capsule, consistent with posterior synechia (Figure 9E). The iridociliary angle remained open.

Within the posterior segment and loosely associated with the detached retina, there were islands and plaques of eosinophilic fibrillary to floccular material, consistent with proteinaceous exudate, alongside small numbers of macrophages, pigment-laden cells and free melanin. The retina exhibited marked, generalised degeneration and was detached from the underlying retinal pigment epithelium (RPE), with residual attachment at the optic disc and ora ciliaris retinae. The RPE underlying the detached neurosensory retina exhibited slight, multifocal hypertrophy (presumed “tombstoning”), supporting chronicity and a true retinal detachment rather than a post-mortem or processing artefact. There was widespread retinal atrophy characterised by almost diffuse loss of ganglion cells, marked loss of both rod and cone photoreceptors. The optic nerve showed moderate, diffuse oedema, with mild vacuolation.

## DISCUSSION

This study demonstrates that iris hyperpigmentation in gannets is associated with clinical and histopathological features consistent with prior intraocular inflammation and with serological evidence of previous HPAI H5Nx exposure. These findings support the interpretation that the widely reported “black-eye” phenotype is due to iris hyperpigmentation and represents a sequela of intraocular inflammation. Histopathology further indicated that iridal hyperpigmentation was characterised by a disruption of normal melanocytic organisation within the iris stroma.

### Ocular morphology and normal variations in gannets

Literature describing ocular morphology in gannets is scarce. Adult and immature birds have pale, forward-facing eyes that provide binocular vision and are protected by a strong, transparent nictitating membrane adapted for plunge-diving (9). Juvenile gannets have a dark iris that lightens with age (3), a finding also described in other seabirds such as Brown Pelicans (*Pelecanus occidentalis*) (7). Histopathological comparison between adult and juvenile eyes showed a loss of a distinct, heavily pigmented anterior border layer of the iris with maturation. A prominent pupillary ruff was observed and represents a normal anatomical feature. However, it may resemble ectropion uveae in the field and, in the absence of other abnormalities, should not be interpreted as evidence of uveitis.

Mean IOP in adult and immature gannets (21.9 ± 4.3 mmHg) was higher than values reported in several bird species, including pigeons and parrots (10). Restraint techniques used during examination (Figure 2) may have contributed to this, as certain methods, such as collars, are known to elevate IOP (11). However, the values recorded here are comparable to those reported in diving birds, including Macaroni Penguins (*Eudyptes chrysolophus*) (21.9 ± 7.05 mmHg) and Rockhopper Penguins (*Eudyptes chrysocome*) (20 ± 5.77 mmHg) (12), and similar to measurements in Ostriches (*Struthio camelus*) (18.8 ± 3.5 mmHg) (13) and Bald Eagles (*Haliaeetus leucocephalus*)) (21.5 ± 1.7 mmHg) (14).This suggests that the relatively higher IOP may reflect species-specific anatomical or physiological adaptations, with restraint-related effects potentially contributing.

### Iridal pigmentary changes and associated ocular inflammation

Our findings suggest that the “black-eye” phenotype in gannets observed during and after the outbreak of HPAI H5N1 (4),(15), is attributable to disorganisation of iridal pigmentation. Among birds exhibiting abnormal iris pigmentation, ranging from mottled to completely black, 77% also displayed clinical signs consistent with uveitis, supporting an association between pigmentary alteration and prior intraocular inflammation. Histopathological examination identified lymphocytic and plasmacytic inflammation within the uveal tract, further supporting this link. Iris hyperpigmentation was associated with disruption of the anterior border layer and melanin disorganisation, including loss of a distinct anterior melanocytic layer and subjective hypertrophy of stromal melanocytes.

Unilateral (n=12) and bilateral (n=19) iris pigmentary changes were observed. The laterality may reflect differences in timing or severity of inflammation; the clinical significance of this laterality remains uncertain.

### Potential mechanisms underlying iris pigmentary changes

Iris colour is influenced by the pigmented epithelium, stromal density and stromal pigment content (16). Chronic prostaglandin-analogue therapy (e.g., latanoprost) in human beings is known to stimulate melanogenesis by upregulating tyrosinase activity, resulting in increased melanin deposition and iris darkening (16). The latanoprost-induced colour change is attributed to thickening of the anterior border layer and increased melanin within both the anterior border layer and stromal melanocytes (17), a pattern that was not observed in the present study. Diffuse iris hyperpigmentation has also been described in canine and feline patients with chronic anterior uveitis, in addition to rubeosis iridis, and is more obvious in eyes with lightly pigmented irides (18). This inflammatory-associated pigmentary change provides a closer clinical parallel to the findings reported here.

In humans, prostaglandin-associated pigmentary changes are considered permanent (19); however, repeat observations of uniquely marked gannets suggest that regression of pigmentation might occur (pers obs; Mackley) although further longitudinal investigation is needed.

Fuchs’ heterochromic cyclitis (FHC) in human beings is a chronic anterior segment inflammatory syndrome of unknown aetiology, accounting for 2–3% of all uveitis cases (20). The condition can lead to characteristic iridal pigmentary changes, most notably heterochromia, in 14% of affected individuals (21). In contrast to the present findings, the affected eye in FHC typically appears lighter due to anterior stromal atrophy, although “reverse heterochromia” may occur when stromal thinning exposes the underlying pigmented epithelium (20). Other human viral infections such as cytomegalovirus (CMV), herpes simplex virus (HSV), and varicella-zoster virus (VZV) can cause iris depigmentation or atrophy, although hyperpigmentation is not reported (22,23). The hyperpigmentation observed in gannets following HPAI infection therefore appears to represent a different inflammatory response.

### Avian influenza and ocular involvement

Ocular involvement is a recognised feature of AIV infection in humans, particularly with H5 and H7 subtypes, where conjunctivitis predominates, although uveitis, retinitis and optic neuritis have been reported (24). Experimental studies demonstrate that multiple AIV subtypes can infect and replicate in human ocular cell types, with the highest viral titres observed for highly pathogenic H7N7 and H5N1 strains, supporting ocular tropism for these viruses (24). In birds, systemic signs of HPAI include high mortality preceded by lethargy, reduced vocalisation, decreased food and water intake, reduced egg production, respiratory signs, neurological abnormalities and diarrhoea (25). Ocular disease is less frequently reported. Corneal opacity associated with endothelial cell loss and keratitis has been described in domestic ducks experimentally infected with H5N1(8), and similar findings were reported in a Dalmatian pelican with serological evidence of infection(26). In naturally infected captive Humboldt penguins (*Spheniscus humboldti)*, HPAI caused severe systemic disease with viral tropism for endothelial and lymphoid tissues, indicating widespread vascular and immune involvement. (27). This endothelial and immune cell tropism provides a plausible mechanistic link to intraocular inflammation and pigmentary changes observed in gannets, although direct viral detection within ocular tissues was not performed in the present study. Evidence of previous ocular inflammation was common, with 36 of 84 examined eyes (from 24 birds) displaying changes consistent with uveitis sequelae. Iris pigmentation abnormalities were positively associated with AIV H5 seropositivity, and 77% of birds with pigmentary changes also exhibited additional signs consistent with intraocular inflammation. Histopathological findings of lymphocytic and plasmacytic uveitis further support this interpretation. In line with the hypothesis proposed by Lane *et al.*(4), these pigmentary changes likely represent sequelae of prior or ongoing inflammatory ocular disease following HPAI infection. A subset of birds exhibiting iridal pigmentary changes tested negative or doubtful for influenza nucleoprotein antibodies on initial screening but subsequently tested positive for H5-specific antibodies. This apparent discordance may reflect differences in antibody kinetics, with nucleoprotein antibodies waning below detectable levels earlier than subtype-specific antibodies. These findings suggest that reliance on nucleoprotein serology alone may underestimate prior HPAI exposure and highlight the value of integrating clinical phenotyping with subtype-specific serology in wildlife disease surveillance. The detection of aqueous flare in one gannet, which was found dead several months later, raises the possibility that clinically apparent active intraocular inflammation may be associated with more severe systemic disease or impaired vision affecting foraging ability.

### Study limitations

This study was conducted under field conditions several years after an HPAI outbreak, which imposed logistical constraints. Concurrent ophthalmic and serological sampling was not possible in all individuals, and examinations were performed in variable natural lighting, which may have reduced sensitivity for detecting subtle abnormalities.

### Conclusions and broader implications

Our findings suggest that striking ocular changes in the gannet represent iris hyperpigmentation, a sequela of inflammatory ocular disease associated with prior HPAI exposure and may provide a visible marker of previous infection in gannets. Neither bird with unaffected eyes and corresponding serology in this study had evidence of a previous HPAI infection, however birds with apparently unaffected eyes have been found to be seropositive for H5 (4, 11) meaning ocular pigmentation cannot currently be used for population level assessments of HPAI exposure.

While current evidence does not demonstrate clear effects on foraging behaviour or reproductive success (11,12), the potential functional consequences for vision remain unknown.

Chronic inflammation may predispose to delayed complications, such as cataract formation, synechiae, or progressive visual impairment, which could have lethal or sublethal consequences, including impaired foraging and survival. While our findings demonstrate a strong association between iris hyperpigmentation and prior HPAI exposure, the reverse is not necessarily true: a normal iris does not guarantee that an individual has not been exposed. Ongoing studies incorporating larger sample sizes and expanded serological testing will help clarify the sensitivity of iris pigmentation as a marker of past infection and identify birds with subclinical or serologically detectable infection without overt ocular signs. More broadly, visible phenotypic alterations such as iris pigmentation may provide an accessible and non-invasive indicator of infectious disease impacts in wildlife populations, offering a valuable tool for long-term monitoring of HPAI and other emerging pathogens.

## AUTHOR CONTRIBUTIONS

Conceptualisation: Chloe Fontaine, Ben Blacklock, David Kayes, Josie Parker, Jude Lane

Data collection: Jude Lane, Jana Jeglinski, Liz Mackley, Kirsty Franklin, Claudia Tapia-Harris, Chloe Fontaine, Ben Blacklock, David Kayes, Josie Parker, Mariana Santos, Adrian Philbey, Liam A. Wilson

Methodology and formal analysis: Hannah Ravenswater, Emma Cunningham, Liz Mackley, Chloe Fontaine

Writing original draft: Chloe Fontaine

Reviewing and editing: Ben Blacklock, David Kayes, Josie Parker, Jude Lane, Mariana Santos, Jana Jeglinski, Liz Mackley, Kirsty Franklin, Claudia Tapia-Harris, Liam A. Wilson, Hannah Ravenswater, Emma Cunningham

## Acknowledgements

We also thank Sir Hew Hamilton-Dalrymple and the Scottish Seabird Centre, North Berwick, for their support and for granting access to Bass Rock, as well as the crew of Seafari for logistical support. We’d like to thank Gidona Goodman, Royal (Dick) School of Veterinary Studies, Edinburgh for Home Office training and support. The authors are additionally grateful to the RSPCA Stapeley Grange Wildlife Centre for providing ocular samples for histopathology. The technical staff in Easter Bush Pathology at the Royal (Dick) School of Veterinary Studies kindly processed samples for histopathology.

## DATA AVAILABILITY STATEMENT

The data that support the findings of this study are available from the corresponding author upon reasonable request.

## ETHICS STATEMENT

Ethical approval for the use of the clinical data was granted by the Ethics Committee of the University of Edinburgh.

